# Noncoding genetic variation in ISPD distinguishes gamecocks from nongame chickens

**DOI:** 10.1101/2023.08.16.553562

**Authors:** Andres Bendesky, Joseph Brew, Kerel X. Francis, Enrique F. Tello Corbetto, Antonio González Ariza, Sergio Nogales Baena, Tsuyoshi Shimmura

## Abstract

Chickens were domesticated >4,000 years ago, probably first for fighting them and only later as a source of food. Fighting chickens, commonly known as gamecocks, continue to be bred throughout the world, but the genetic relationships among geographically diverse gamecocks and with nongame chickens are not known. Here, we sequenced the genomes of 44 geographically diverse gamecocks and of 62 nongame chickens representing a variety of breeds. We combined these sequences with published genomes to generate the most diverse chicken genomes dataset yet assembled, at 307 samples. We found that gamecocks do not form a homogeneous group, yet they share genetic similarities that distinguish them from nongame chickens. Such similarities are likely the result of a common origin before their local diversification into, or mixing with, nongame chickens. Particularly noteworthy is a variant in an intron of ISPD, an extreme outlier present at a frequency of 90% in gamecocks but only 4% in nongame chickens. The ISPD locus has the strongest signal of selection in gamecocks, suggesting it is important for fighting performance. Because ISPD variants that are highly prevalent in gamecocks are still segregating in nongame chickens, selective breeding may help reduce its frequency in farm conditions in which aggression is not a desired trait. Altogether, our work provides genomic resources for agricultural genetics, uncovers a common origin for gamecocks from around the world and what distinguishes them genetically from chickens bred for purposes other than fighting, and points to ISPD as the most important locus related to fighting performance.

## Introduction

Chickens were one of the first animals to be domesticated, ∼4,000-10,000 years ago in South Asia, from the Red junglefowl (*Gallus gallus*)^1–6^. They are now the most abundant land vertebrate, with a population size of over 33 billion and more than 25 billion chickens produced every year^7^. Today, chickens are by and large used for egg and meat consumption. Archaeologists propose, however, that chickens may have been first domesticated as gamecocks for fighting purposes and that this use drove their spread throughout the world^2,8–10^. Evidence of their use for fights dates to at least 2,500-2,700 BC in the Indus Valley^2^ and China^9^.

Organized chicken fights are still common and ingrained in cultures throughout the world^11,12^. The chickens used in fights are distinct from those bred as a source of food; instead they are selectively bred for fighting. Some gamecocks are bred as defined breeds, such as the Gallo Combatiente Español, the Gallo Navajero Peruano, the French Combattant du Nord, and varieties of Japanese Shamo, whereas others are more mixed. In some places, gamecocks are no longer used in fights but are still actively produced to maintain historical breeds, which can trace their origins to hundreds of years^13–15^.

Gamecock breeds vary in many traits but share a key behavioral feature^13^. Some breeds are specialized for endurance in lengthy, hours-long fights, whereas others fight much shorter fights, aided by different kinds of sharp instruments attached to their legs by people^16^. Gamecocks also vary drastically in their anatomy: there is a 350% range in their body weight, from the compact 2-kg Red Cubans^13^ to the tall 7-kg Shamo^17^. Thus, gamecocks vary in endurance, fighting style, body shape, and mass. By contrast, a behavioral feature shared by gamecocks, which sets them apart from nongame chickens, is their so-called “gameness” —a strong tendency to engage in a fight with another cock, rather than to flee, even after injury^18^. Breeders directly select for gameness in behavioral tests before sanctioned fights, and indirectly by selectively breeding cocks (and their first-degree relatives) that have won fights^13,15,18–20^. The ultimate purpose of this strong selection for gameness and fight performance is to produce game chickens that will win fights.

There is consensus among gamecock breeders that nongame chicken are much less willing to engage in and persist in a fight than gamecocks (ref. ^15^ and personal communications from multiple breeders in Mexico, Peru, and Puerto Rico). Nevertheless, there are no scientific reports of the heritability or genetic bases of gameness. A recent genome-wide scan of selection in Chinese gamecocks identified multiple loci with low heterozygosity and that these loci genetically distinguish them from Chinese nongame chickens^9^. However, genetic scans of selection in gamecocks outside China have not been performed, so it is unknown if the same or different loci are involved in gamecocks globally. Furthermore, how gamecocks from around the world are related to each other and to local and global nongame chickens is not known.

Although the history of chicken domestication suggests an ancient and common origin of gamecocks, it is possible that different gamecock breeds arose independently, from local chicken populations. Here, we set out to discover the genetic relationships among gamecocks from around the world, and to that end also assembled a phenotypically and geographically diverse set of samples from chickens bred for purposes other than fighting. We then searched for genetic similarities among gamecocks of different breeds for genetic signatures of selection that would point to possible causes of their fighting performance.

## Results

To study the genetics of gamecocks, we began by collecting DNA samples of 44 fighting chickens from around the world and sequencing their genome to ∼12× coverage. After combining them with four other published gamecock sequenced genomes, this dataset includes chickens representing 15 countries and 29 breeds (**Table 1**). As a comparison group of chickens bred for purposes other than fighting, we further sequenced 62 nongame chickens and used 181 additional published nongame chicken samples (**Suppl. Table 1**). Together, the set includes samples from 12 countries and 108 recognized chicken breeds, or chickens bred in a particular location but without a particular breed name. Many of these breeds had never been sequenced. This set of samples thus constitutes the most diverse collection of chicken genomes sequenced at relatively high coverage.

**Table 1.** Gamecock genomes sequenced and analyzed in this study.

| Sample origin | Breed | Number of samples | Source |
| --- | --- | --- | --- |
| Bali | Balinese | 1 | This study |
| Brazil | Shamo | 1 | This study |
| China | Luxi game | 1 | Ref.<sup>21</sup> |
| China | Xishuangbanna game | 3 | Ref.<sup>22</sup> |
| Colombia | Pico | 1 | This study |
| France | Combattant du Nord | 1 | This study |
| Japan | Koshamo-Fukushima | 1 | This study |
| Japan | Nankinshamo-Hiroshima | 1 | This study |
| Japan | Oshamo | 14 | This study |
| Japan | Tauchi | 1 | This study |
| Japan | Yakido-Fukushima | 1 | This study |
| Japan | Yakido-Hyogo | 1 | This study |
| Japan | Yakido-Mie | 1 | This study |
| Malaysia | Omkoï | 1 | This study |
| Mexico | Hatch | 1 | This study |
| Mexico | Mixed | 1 | This study |
| Mexico | Navajero | 1 | This study |
| Mexico | Sweater | 1 | This study |
| Pakistan | Asil | 1 | This study |
| Peru | Navajero Peruano | 1 | This study |
| Peru | Pico | 2 | This study |
| Puerto Rico | Pico | 1 | This study |
| Spain | Combatiente Español | 1 | This study |
| Spain | Combatiente Canario | 1 | This study |
| Thailand | Nakhon Sawan | 1 | This study |
| USA | Black Breasted Red | 1 | This study |
| USA | Blue Sumatra | 1 | This study |
| USA | Dark Aseel | 1 | This study |
| USA | Sumatra | 1 | This study |
| USA | Wheaton Aseel | 1 | This study |
| Vietnam | Saigon | 1 | This study |

### Gamecocks do not constitute a homogeneous group

To determine whether gamecocks from around the world are more closely related to each other than to chickens not bred for fighting, we built a phylogenetic tree based on whole genome data of the 48 gamecock samples and the 243 nongame domesticated chickens (**Figure 1** and **Suppl. Figure 1**). To build this tree, we also included published genomes of Red junglefowl (*Gallus gallus*), from which the chicken was domesticated^1,5,6,23^, as well as from the closely-related Ceylon junglefowl (*Gallus lafayetii*), the Grey junglefowl (*Gallus sonneratii*), and, as an outgroup, the more distantly related Green junglefowl (*Gallus varius*). As expected, the Green junglefowl formed an outgroup to all other samples. The Ceylon and Grey junglefowls appear as sister species to each other and are more closely related to Red junglefowls than to Green junglefowls, in agreement with previous observations^1,23^. All chickens, including gamecocks, are included in a monophyletic branch that is distinct but closest to the Red junglefowls, consistent with chickens being domesticated from the Red junglefowl.

**Figure 1.**
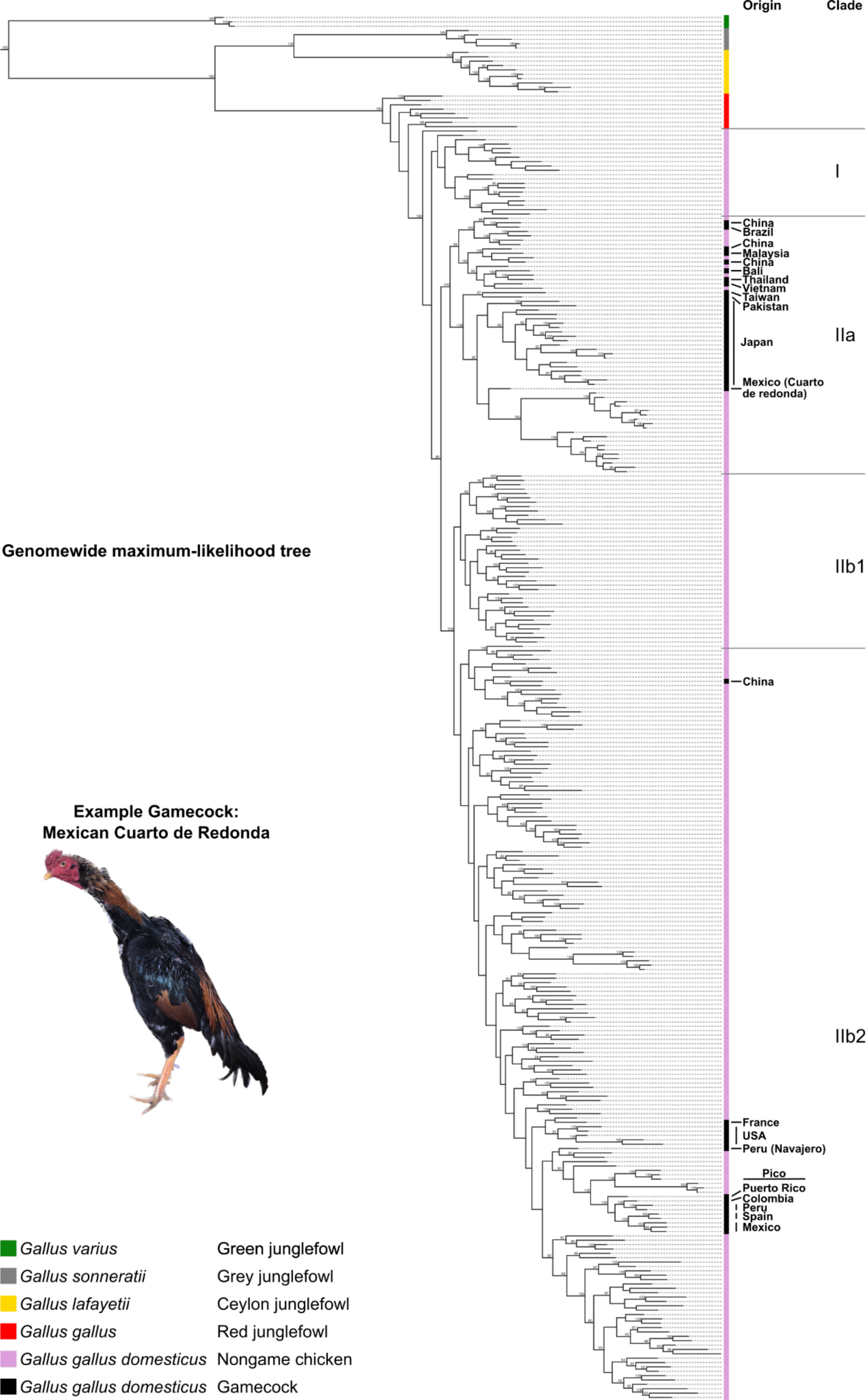
Phylogenetic tree of *Gallus* including chickens. Maximum-likelihood phylogenetic tree based on whole-genome data, including all species in the junglefowl (*Gallus*) genus, as well as gamecocks and nongame chickens from around the world. The geographic origin of gamecock breeds is highlighted on the right. Bootstrap support values ≥90 are highlighted on the branches. Suppl. Figure 1 is a zoomable and text-search friendly version of the figure showing the location and name of all samples.

Chickens separate into two major clades: I and II. Clade I contains exclusively nongame chickens from China and Tibet. Clade II is further divided into IIa and IIb. IIa contains almost exclusively Asian nongame and game chickens as well as a gamecock sample from Mexico and one from Brazil; the breeders of these Latin American gamecocks acknowledged used Asian gamecocks to produce these individuals, likely explaining their phylogenetic placement. Clade IIb is subdivided into two further clades. Clade IIb1 is composed exclusively of non-game chickens from China. By contrast, Clade IIb2 is the most diverse group, including samples from most locations. All the European, African, and New World game and nongame chickens (except the Mexican and Brazilian ones noted above) are contained in this clade. This clade also contains chickens from East Asia and Iran. Together, these results show that chicken genetic relatedness is broadly geographically-structured, but with some exceptions likely originating from recent mixing with animal imports from other locations.

Notably, gamecock samples do not all cluster together but rather are interspersed with nongame chickens, often with individuals from the same geographic region (**Figure 1** and **Suppl. Figure 1**). The lack of a single gamecock cluster suggests that gamecocks have independent local origins or a common origin followed by mixing with local populations.

### Phylogenetic patterns reflect historical events

The patterns of relationships among gamecocks appear to parallel their history. Fighting chickens from North America, South America, and the Caribbean are most closely related to the Combatiente Español and Combatiente Canario gamecock breeds from Spain, consistent with the introduction of gamecocks into the New World by the Spanish^8,14^ (**Suppl. Figure 1**). These Hispanic and New World gamecocks are most closely related to the Leghorn (Livornese) chickens that originated in Tuscany, possibly related to an introduction of gamecocks into Spain by the Romans^8,13^. The close relationship between Hispanic gamecocks and Italian nongame chickens indicates these breeds share more ancestry than other breeds.

### Gamecocks share genetic similarities

The genome-wide phylogeny indicated that gamecocks do not form a homogeneous group. Nonetheless, gamecocks may still share genetic ancestry distinct from nongame chickens in a fraction of their genome, indicative of a common origin. To test this possibility, we began by performing principal component analysis (PCA) of genome-wide genetic variation. The first three principal components cleanly separated samples by species (**Figure 2a**). Gamecocks clustered closely with nongame chickens and with Red junglefowl in the first two PCs (**Figure 2a**), suggesting no large-scale hybridization of any of these *Gallus gallus* samples with other *Gallus* species, in agreement with the genome-wide phylogeny. PC 5 largely separated *Gallus gallus* into domesticated and wild individuals (**Figure 2b** and see **Suppl. Figure 2a**). In turn, PC 12 largely separated domesticated chickens into game and nongame (**Figure 2c** and see **Suppl. Figure 2b**). Remarkably, this PC also separated Japanese chickens into game and nongame (**Figure 2d** and see **Suppl. Figure 2c**), even though these two groups are otherwise closely related and form sister branches in the genome-wide phylogeny (**Suppl. Figure 1**). These results indicate that gamecocks from around the world share genetic similarities.

**Figure 2.**
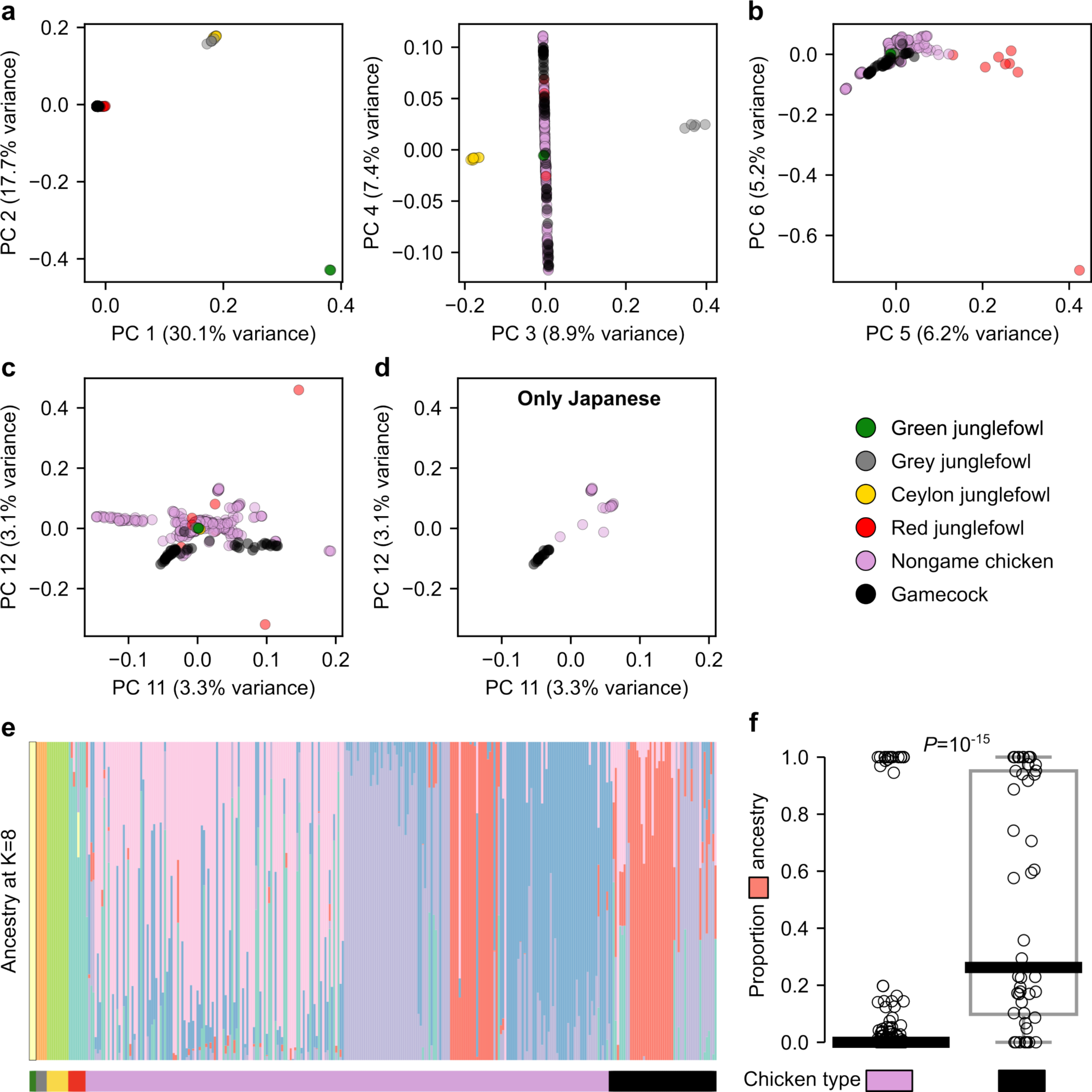
*Gallus* genetic structure a-d, Principal Component Analysis of genetic variation in *Gallus* samples. **a,** PC 1-3 separate samples by species. **b,** PC 5 separates *Gallus gallus* into Red junglefowl and domesticated chickens. **c,** PC 12 mostly separates chickens into gamecocks and nongame chickens. **d,** Same as **c** but showing Japanese samples exclusively. **e,** ADMIXTURE analysis at K=8 (which had the lowest cross-validation error, see Suppl. Figure 3a). **f,** Proportion of “salmon-colored” ancestry from **e** in nongame chickens (n=243) and gamecocks (n=48). *P*-value by Mann-Whitney U test.

Consistent with these findings, an ADMIXTURE analysis to characterize genetic ancestry components revealed that gamecocks have significantly (*P*=10^-15^ by a Mann-Whitney U test) more of a particular ancestry (median of 26%) than do nongame domesticated chickens (median of 0%) (**Figure 2e,f** and **Suppl. Figure 3a**). This “game ancestry” was not driven by the Japanese samples, which constitute 15% of the dataset, as the game ancestry remains significantly (*P*=10^-10^) more common in non-Japanese gamecocks than in non-Japanese nongame chickens (**Suppl. Figure 3b**). The game ancestry does not appear to correspond to that of closely related *Gallus* species, suggesting that it is unlikely the product of incomplete lineage sorting or gene flow from those species (**Figure 2e**). Thus, although gamecocks do not form a homogeneous phylogenetic group, they share a significant component of their ancestry. Even though gamecocks had significantly more of this game ancestry, some gamecocks lacked it and some nongame chickens had substantial game ancestry, suggesting that only a fraction of this ancestry might be relevant for fighting-related traits. A parsimonious explanation for these observations is that gamecocks from around the world had a common origin, which was followed by local diversification into or local mixing with nongame chickens, but that some of their original game ancestry —perhaps that which is most relevant for fighting performance— was maintained through selective breeding, despite local mixing.

### Gamecocks share genetic variants under selection

To identify genetic loci shared by gamecocks, we performed a genome-wide association study (GWAS) comparing gamecocks to nongame chickens, using hierarchical clustering to control for population structure^24^ (**Figure 3a** and **Suppl. Figure. 4b**). We found a particularly prominent peak in Chromosome 2, near the genes *alkylglycerol monooxygenase* (AGMO), *mesenchyme homeobox 2* (MEOX2), an uncharacterized RNA with an open reading frame, and reaching the zenith in an intron of *isoprenoid synthase domain containing* (ISPD) (**Figure 3b**). Intriguingly, this locus was recently identified in an independent study as the most differentiated (highest *F*_ST_) genomic region between Chinese gamecocks and Chinese nongame chickens^9^. The GWAS peak we identified is not driven by Chinese chicken samples, since they only constitute 8% of our game samples and none of our nongame samples, and the peak remains the top hit when excluding Chinese chickens (**Suppl. Figure 4a,c**). Thus, this Chromosome 2 locus is the genetic region that most distinguishes gamecocks from nongame chickens worldwide.

**Figure 3.**
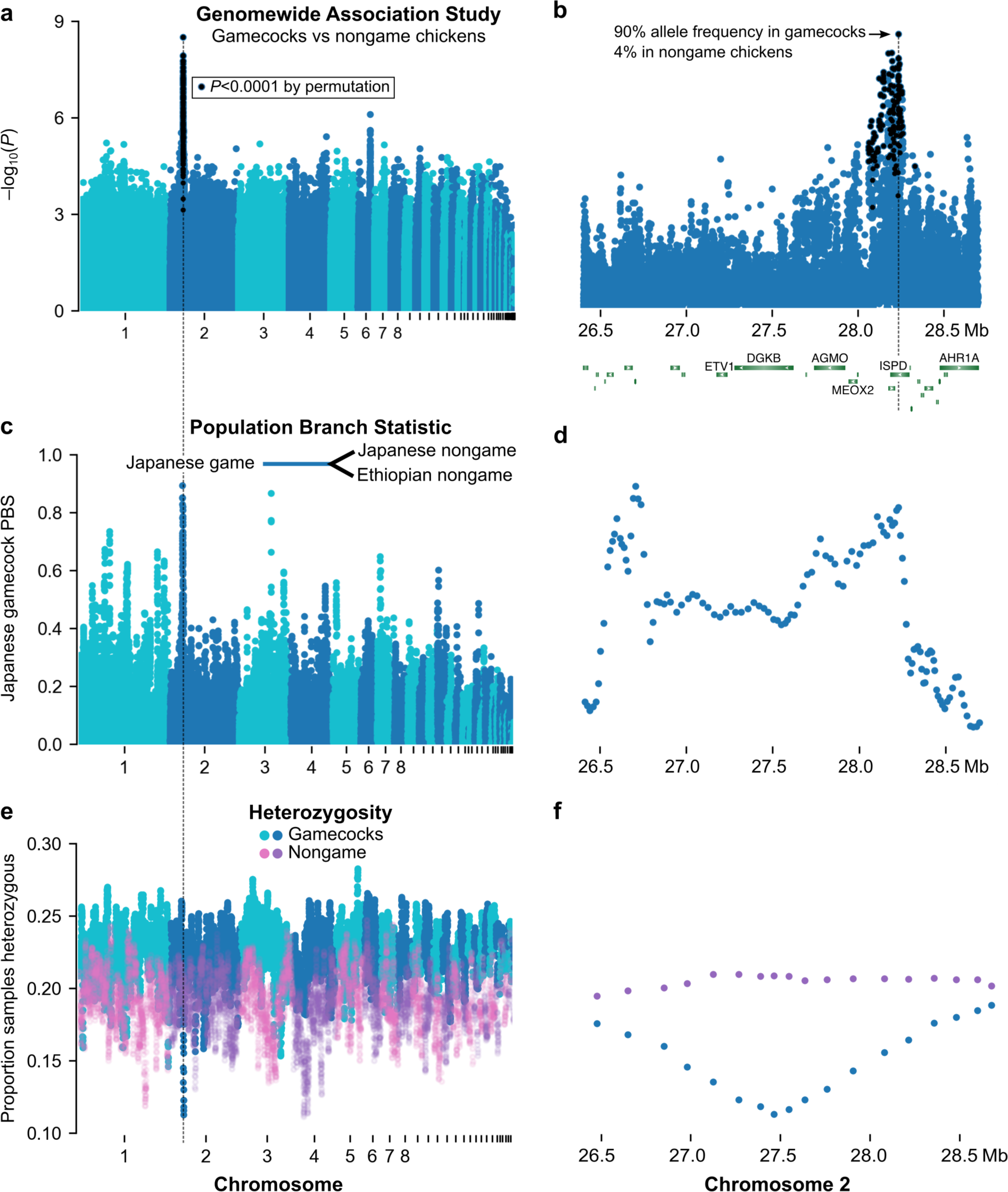
Genetic loci that distinguish gamecocks from nongame chickens are under selection. **a,b,** Genome-wide Association Study comparing gamecocks (n=48) to nongame chickens (n=62) and zoom of top hit. *P*-values on the y axis using genomic control. Black denotes variants with *P*<10^−4^ by permutation; no variants outside chromosome 2 surpassed that permutation threshold. **c,d,** Windowed Population Branch Statistic comparing Japanese gamecocks (n=14) to Japanese nongame chickens (n=19) and Ethiopian nongame chickens (n=34). The Japanese gamecock branch statistic is shown on the y axis. **e,f,** Windowed heterozygosity in gamecocks (n=48) and nongame chickens (n=62).

**Figure 4.**
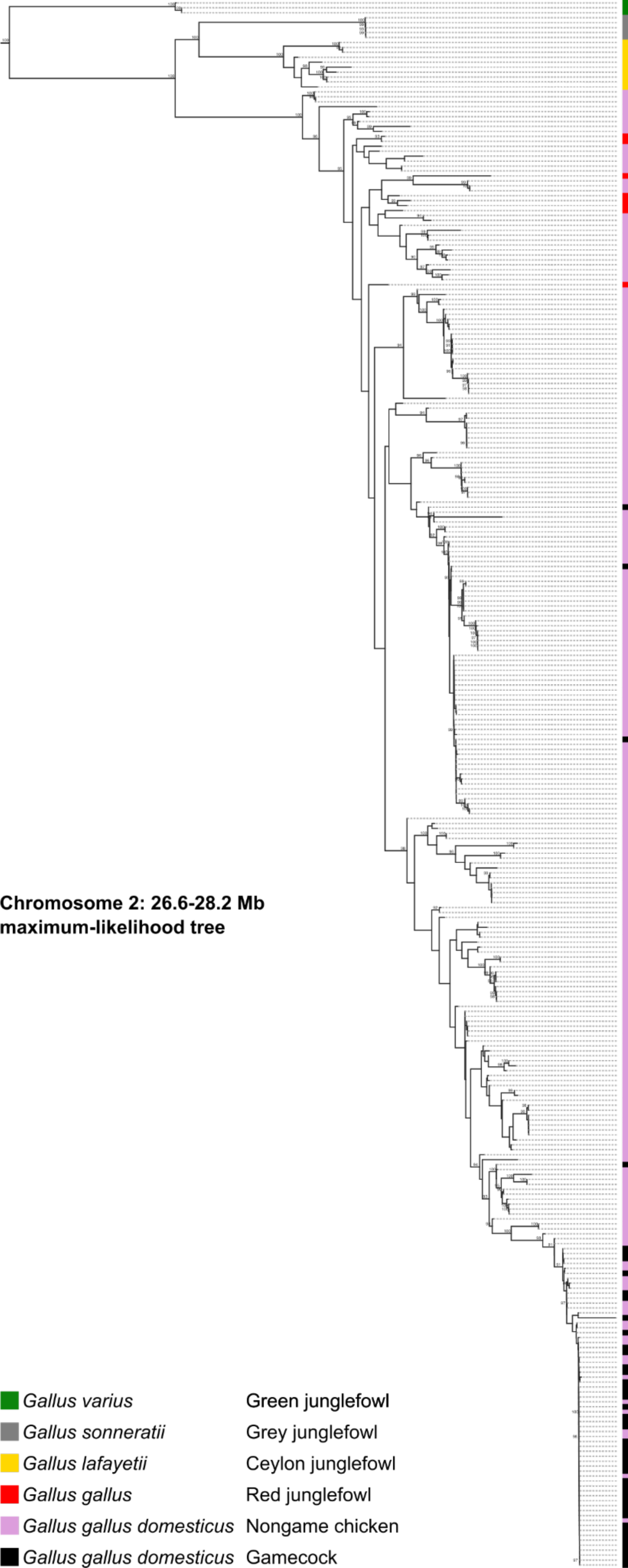
Phylogenetic tree of Chromosome 2 locus that distinguishes gamecocks from nongame chickens. Maximum-likelihood phylogenetic tree of Chromosome 2 locus (26.6–28.2 Mb), including all species in the junglefowl (*Gallus*) genus, as well as gamecocks and nongame chickens from around the world. Bootstrap support values ≥90 are highlighted on the branches. Suppl. Figure 5 is a zoomable and text-search friendly version of the figure showing the location and name of all samples.

The most significantly (Genomic Control *P*=3×10^-9^; permutation *P*<10^-4^) associated variant (rs735074467), at position 28,236,213 in Chromosome 2 within an intron of the ISPD gene, is present at a frequency of 4% in nongame chickens but 90% in gamecocks (**Figure 3b**). By contrast, the ISPD missense variant Arg84Lys that was previously identified as a likely causal variant in Chinese gamecocks^9^ is present at a frequency of 3% in nongame chickens but only 37% in gamecocks and it does not reach genome-wide significance (Genomic Control *P*=1.7×10^-3^; permutation *P*=1) in our GWAS. Thus, noncoding variation in ISPD is a more likely genetic cause of differences between gamecocks and nongame chickens worldwide.

If gamecocks had a common origin and later mixed with nongame chickens, gamecock ancestry could be diluted randomly, in which case the Chromosome 2 gamecock-associated variants remain in most gamecocks by chance. Alternatively, the Chromosome 2 locus variants might persist in gamecocks through the continued effects of selective breeding. Arguing in favor of the selective breeding hypothesis, a population-branch statistic (PBS) analysis indicates that the Chromosome 2 locus is the part of the genome most differentiated in Japanese gamecocks compared to closely related Japanese nongame chickens and to Ethiopian nongame chickens as a more distant control population, consistent with selection at the locus (**Figure 3c,d**). In line with the PBS results, the Chromosome 2 locus (the 1.6 Mb plateau with high PBS) has 38% lower nucleotide diversity in gamecocks (0.1% per site) than in nongame chickens (0.16%); and, notably, the locus has the lowest heterozygosity of the whole genome in gamecocks from throughout the world, but has unremarkable heterozygosity in nongame chickens, consistent with selection in gamecocks (**Figure 3e,f**). This region also has unusually low heterozygosity in Chinese gamecocks^9^. Altogether, our results indicate that the Chromosome 2 locus is the region that most strongly distinguishes gamecocks from nongame chickens, likely because of the effects of selective breeding.

Because gamecocks share genetic ancestry, it is likely that the Chromosome 2 locus of gamecocks has a common origin, i.e., this region is identical by descent among them. Alternatively, the locus evolved independently and convergently in gamecocks. Consistent with a common origin, 92% (44/48) of gamecocks, including those from clades IIa and IIb1, cluster together in a branch of a phylogenetic tree of the 1.6-Mb Chromosome 2 PBS locus (**Figure 4** and **Suppl. Figure 5**). By contrast, only 9% (21/243) of nongame chickens are in this grouping (*P*<10^-4^ by Fisher’s exact test). Very similar phylogentic patterns are observed in a narrower 200-kb region encompassing the GWAS peak (**Suppl. Figure 6**). In agreement with the phylogenetic clustering, the gamecock and nongame haplotypes are very different (**Suppl. Figure 7**). Thus, although gamecocks differ substantially from each other genome-wide (**Figure 1**), they are very similar at the Chromosome 2 locus. All junglefowl samples follow the species tree at this locus, suggesting that this gamecock haplotype is not the result of introgression from another junglefowl species. Thus, the locus that is most associated with gamecocks has a common origin across gamecocks.

## Discussion

We assembled the most diverse panel of gamecocks and nongame chickens, of the Red junglefowl (the wild ancestor of the domesticated chicken), and of the other three species in the junglefowl (*Gallus*) genus. Through the analysis of whole genomes, we found that gamecocks from around the world (Bali, Brazil, China, Colombia, France, Japan, Malaysia, Mexico, Pakistan, Peru, Puerto Rico, Spain, Taiwan, Thailand, USA, and Vietnam) share common ancestry that is largely absent from nongame chickens. This pattern is likely the result of positive selection for high aggression in gamecocks and against aggression in nongame chickens, in which it is an undesirable trait^25^.

Genome-wide, however, gamecocks do not form a homogeneous group. One possibility is that chickens were repeatedly and independently selectively bred for fighting around the world and parallel selection on standing variation increased the frequency of “gamecock ancestry” in geographically disperse chickens. Alternatively, gamecocks could have had a common origin and later mixed with local nongame populations as they dispersed through the world. The mixing with local populations could have diluted the “gamecock ancestry” through the random effects of recombination and the independent assortment of chromosomes at meiosis, such that specific loci would now be enriched in gamecocks only by chance. However, we find a locus on Chromosome 2, peaking at the gene ISPD, that strongly differs in variant frequencies between gamecocks and nongame chickens from around the world. This region had been found to distinguish Chinese gamecocks from Chinese nongame chickens^9^; our results now show that this locus is the most distinguishing genomic locus for gamecocks worldwide and provides a much finer resolution. Moreover, we find evidence for this locus being under selection in gamecocks, as it is particularly differentiated in gamecocks relative to the rest of the genome and shows the lowest levels of heterozygosity in the genomes of gamecocks but not in nongame chickens. Consistent with a common gamecock origin, this locus, in contrast to the rest of the genome, is very similar in most gamecocks.

Because gameness and fighting ability distinguishes gamecocks from nongame chickens more than body size or other anatomical features, it is likely that the Chromosome 2 locus contributes to the behavior of gamecocks. It is possible, however, that it affects other traits that modulate fighting performance, such as athleticism. Previous reports focusing on Chinese gamecocks pointed to a missense (Arg84Lys) variant in ISPD within the Chromosome 2 locus as likely causal and that this variant might affect muscle traits^9,26^. However, our results, which encompass more geographically and genetically diverse gamecock and nongame chicken samples, as well as a larger sample size, indicates the ISPD missense variant is not significantly more frequent in gamecocks. Instead, our results point to an intronic variant in ISPD as the most pronounced genetic distinction between gamecocks and nongame chickens. Although genetic variation in and near ISPD may act by affecting other genes, ISPD itself is an appealing candidate not only because it’s the gene closest to the peak of association, but also because ISPD is involved in axon guidance during the wiring of the central nervous system and in the development of the cerebral cortex^27^. Regulatory variation at ISPD may therefore affect fighting performance through behavioral effects rather than, or in addition to, its effects on muscle.

Altogether, this work characterizes the commonalities and differences among the genomes of gamecocks and points to a locus in Chromosome 2 as a genetic factor under selection that almost perfectly distinguishes gamecocks from other chickens. Because genetic variants associated with gamecocks are still segregating in nongame chickens, it might be possible to use these markers to select chickens with reduced aggression for breeding under conditions in which aggression is not a desirable trait.

## Methods

### Sample acquisition and whole genome sequencing

Animal protocols were approved by the Institutional Animal Care and Use Committee of Columbia University. We obtained blood samples after informed consent from breeders. We drew blood from the wing vein, deposited in Whatman 903 cards, let dry completely and then stored in a ziploc bag at -20 °C until DNA extraction. We extracted DNA from a 3.2-mm circular punch of the blood card by digesting in proteinase K with SDS and Tween 20 followed by Omega Magbind Blood and Tissue kit. We then created Tn5-tagmented, 9-PCR cycles amplified, whole-genome sequencing libraries^28^ of an average insert size of 300-400 bp. We sequenced libraries with 2×150bp reads in an Illumina Novaseq to a median depth of 12.2× (minimum 8×) after removing duplicate reads (see below).

Additionally, we downloaded sequencing data of 32 individuals with >9× coverage from the Chicken SD database, http://bigd.big.ac.cn/chickensd/. We also downloaded resequencing data of 300 samples with >8× coverage across all four *Gallus* species and 59 named chicken breeds from 12 NCBI BioProjects: PRJDB4092^21^, PRJEB39275^29^, PRJNA202483^30^, PRJNA232548^21^, PRJNA241474^22^, PRJNA306389^31^, PRJNA432200^5^, PRJNA482210^32^, PRJNA524911^33^, PRJNA547951^34^, PRJNA552722^35^, PRJNA597842^36^, and PRJNA659069^37^.

### Variant calling and filtering

We followed GATK best practices for mapping and cleaning up short read sequence data efficiently^38^: 1) we converted fastq files to uBAMs using picard FastqToSam^39^; 2) we marked adapters (NEXTERA_V2) using picard MarkIlluminaAdapters; 3) we aligned reads to the bGalGal1.mat.broiler.GRCg7b (GCF_016699485.2, part of the Vertebrate Genomes Project^40^) *Gallus gallus* reference genome by piping picard SamToFastq output to bwa mem2^41^ and piped the output to picard MergeBamAlignment; and 4) we merged alignments of the same individual coming from multiple sequencing runs and marked duplicate reads using picard MarkDuplicates.

We called variants using bcftools mpileup^42^ and then filtered individual calls and then sites from all individuals using bcftools according to the following criteria. For individual calls, we included only variants >3 bp from an indel, that had an MQ>=50.0 and MQ0F<=0.1, excluded indels and variants with more than 2 alleles, included only autosomal variants whose DP was <1.65× the average autosomal DP for that sample^43^, excluded variants with DP<4 GQ<30, that were het yet AD<2 for either allele, that were het yet the AD ratio of one allele to the other was <0.25 or >0.75. For sites, we excluded those that had missing genotypes (according to the filtering criteria above) in more than 20% of samples. We also excluded sites that fell in poor mapability regions according to a snpable^44^ mask of the bGalGal1 genome ran with the following parameters: kmer length k=100, map with bwa aln -t 16 -R 1000000 -O 3 -E 3, stringency r=0.5.

### Sample filtering and quality control

We quality-checked the sequencing quality of each sample using fastp^45^. We excluded downloaded sample SRR1217514 because over 99% of the reads had low quality (average read quality score <Q15). We further removed samples from the downloaded dataset that did not have >7.5× coverage after duplicates had been removed. The median depth of the remaining downloaded samples was 12.6×. We also ensured that no two samples were identical using picard crosscheck fingerprint, nor more closely related than first cousins using ngsRelateV2^46^, nor showed signs of cross-contamination based on variant allele frequencies >20% across the mitochondrial genome. Finally, we removed samples that had more than 30% genotypes missing after all the site and genotype filters above had been applied. An additional published Red junglefowl sample (SRR1217528) was removed because it clustered with domesticated chickens rather than with all other Red junglefowl in both the genome-wide phylogenetic tree and in PCA, suggesting this was a Red junglefowl–chicken hybrid or the result of sample contamination.

### Phylogenetic analyses

For the genome-wide phylogenetic tree, we sampled ∼0.1% of the genome by selecting 1,000 non-overlapping 10-kb regions randomly distributed throughout the genome except the Z and W chromosomes, and generated multi-fasta files from a bcf file containing invariant sites. Using this multi-fasta file, we generated a genome-wide maximum likelihood tree using IQTree v2.0.3^47–49^ with the following parameters: -t PARS -ninit 2 -bb 1000 -nm 1000 -m TEST (to find the best-fit substitution model for the data, which was HKY+F+I). For the phylogenetic tree of the Chromosome 2 PBS (26.6–28.2 Mb) and GWAS (28,061,838–28,260,724 bp) peaks, we ran IQTree on fasta files including invariant sites of the regions using the following parameters: -t

~~~
PARS -ninit 2 -bb 1000 -nm 5000 -m HKY+F+I. We plotted the trees using toytree 2.0.5^50^.
~~~

### Principal component analysis and ADMIXTURE

For genetic principal component analysis, we used PLINK v1.90b6.21^51,52^ on autosomal variants with a minimum allele count of 1 with the following parameters for linkage disequilibrium pruning of variants --indep-pairwise 50 kb window size 10 kb step size 0.2 r^2^ threshold. We plotted the results using seaborn^53^. We ran ADMIXTURE v1.3^54^ at K=1-8 using the plink linkage-disequilibrium pruned variants at 0.1 r^2^ threshold and present the results for K=7 because it had the lowest cross-validation error.

### Genome-wide association analysis

We performed genome-wide association analysis (GWAS) comparing gamecocks —using the samples we collected specifically for this study (plus the 4 published Chinese gamecock samples) and thus were confident on their breeding for fighting (cases, n=48)— to nongame chickens (controls, n=62). We used the 9,023,221 autosomal variants at a minimum allele frequency of 5%. To control for population stratification, we first clustered the samples using complete linkage agglomerative clustering based on pairwise identity-by-state distance, using PLINK v1.90b6.21. Both K=4 and K=5 led to the lowest genomic inflation compared to K=2,3,6,7,8,9,10. However, K=4 included more samples in clusters than K=5 so we performed association testing using K=4, accounting for clusters, and calculated statistical significance using genomic control.

~~~
--cluster --K 4 --allow-extra-chr --allow-no-sex --out out
--mh --within out.cluster2 --adjust --allow-extra-chr --allow-no-sex --out aac
~~~

We then performed 10,000 permutations, also accounting for clusters with K=4 and used a permutation-based significance threshold of *P*<10^-4^.

### Population branch statistic, heterozygosity, and nucleotide diversity

We calculated population branch statistic (PBS)^55^, with the focal population being Japanese gamecocks, and the Japanese nongame chickens and Ethiopian nongame chickens as the contrasts. We used variants with a minimum allele count of 1. PBS was calculated in windows of 1,000 variants with 200 step size, using scikit-allel v.1.3.5^56^.

Heterozygosity was calculated separately in gamecocks and nongame chickens (as described in GWAS section), in windows of 10,000 variants and 1,000 step size, using scikit-allel v.1.3.5.

Nucleotide diversity within the PBS peak region (Chromosome 2: 26.6–28.2 Mb) was calculated using pixy v1.2.7beta1^57,58^ on a vcf file that included invariant sites, separately in gamecocks and nongame chickens (as described in GWAS section).

### Phasing

To plot haplotypes, we phased genotypes using SHAPEIT5 v5.1.0^59^ phase_common using default parameters.

## Author contributions

AB and TS conceived the project and collected samples. EFTC, AGA, and SNB provided samples. KXF extracted DNA and generated sequencing libraries. JB identified suitable public datasets and performed preliminary analyses. AB analyzed the data and wrote the manuscript.

## Acknowledgements

Federación Gallística Canaria, Unión de Criadores de Gallos Combatiente Español, Cackle Hatchery, and chicken breeders around the world provided samples. Víctor Pérez assisted in the field. Natalie Niepoth and Carla Hoge helped collect samples. Young Mi Kwon helped with analyses. Molly Przeworski provided comments on the manuscript. **Funding:** National Institutes of Health grant R35GM143051, Searle Scholarship, Klingenstein-Simons Fellowship in Neuroscience, and Sloan Fellowship in Neuroscience to AB. JSPS Grant-in-Aid for Scientific Research (B) 21H00339, and JST FOREST Program JPMJFR211D, Japan, to TS.

## Data availability

Sequence data is available at NCBI Sequence Read Archive BioProject ID PRJNA973554.

---

**Suppl. Figure 1.**
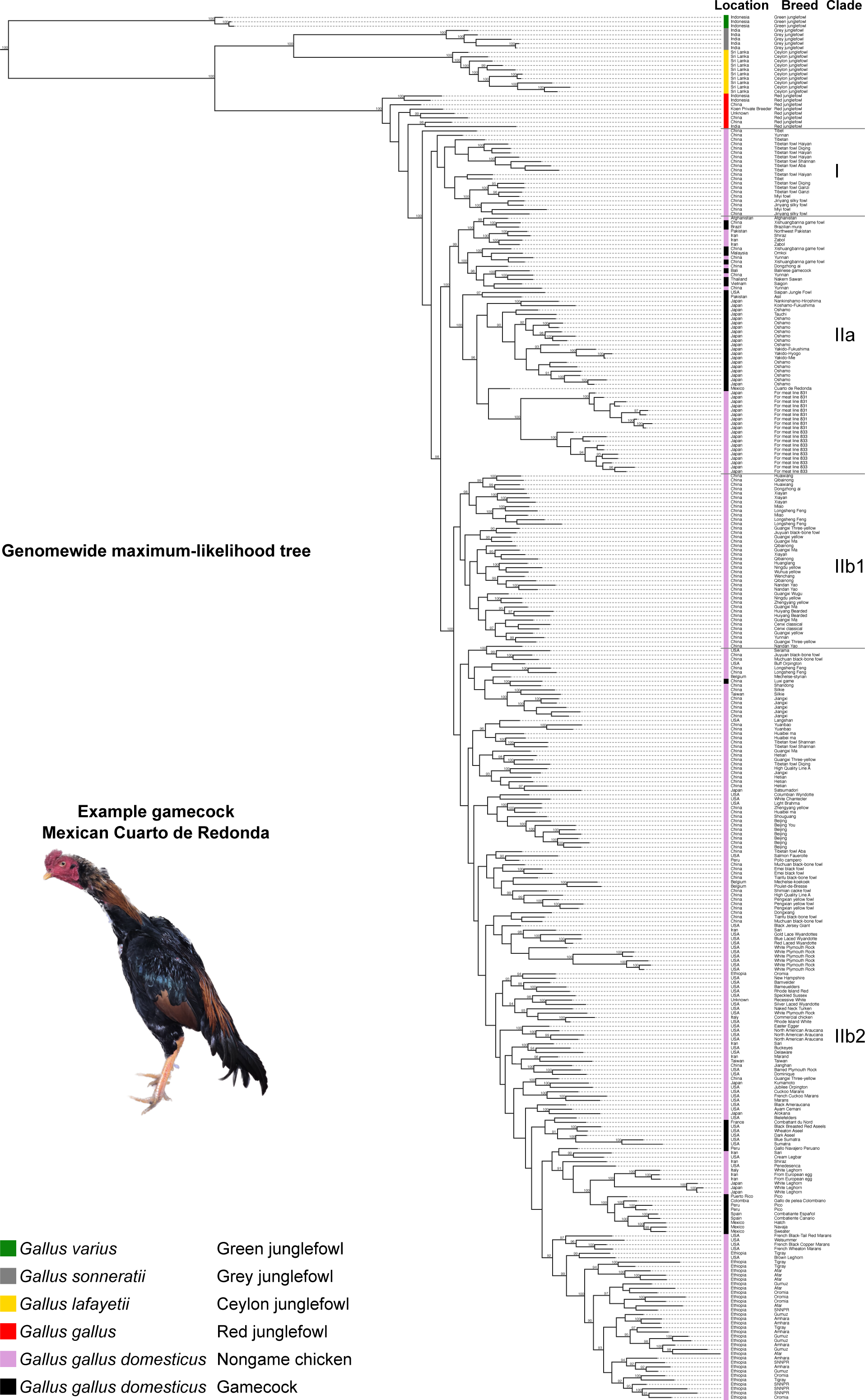
Genome-wide phylogenetic tree of *Gallus* including chickens. Maximum-likelihood phylogenetic tree based on whole-genome data, including all species in the junglefowl (*Gallus*) genus, as well as gamecocks and nongame chickens from around the world. The name and geographic origin of the samples is presented on the right. Bootstrap support values ≥90 are highlighted on the branches.

**Suppl. Figure 2.**
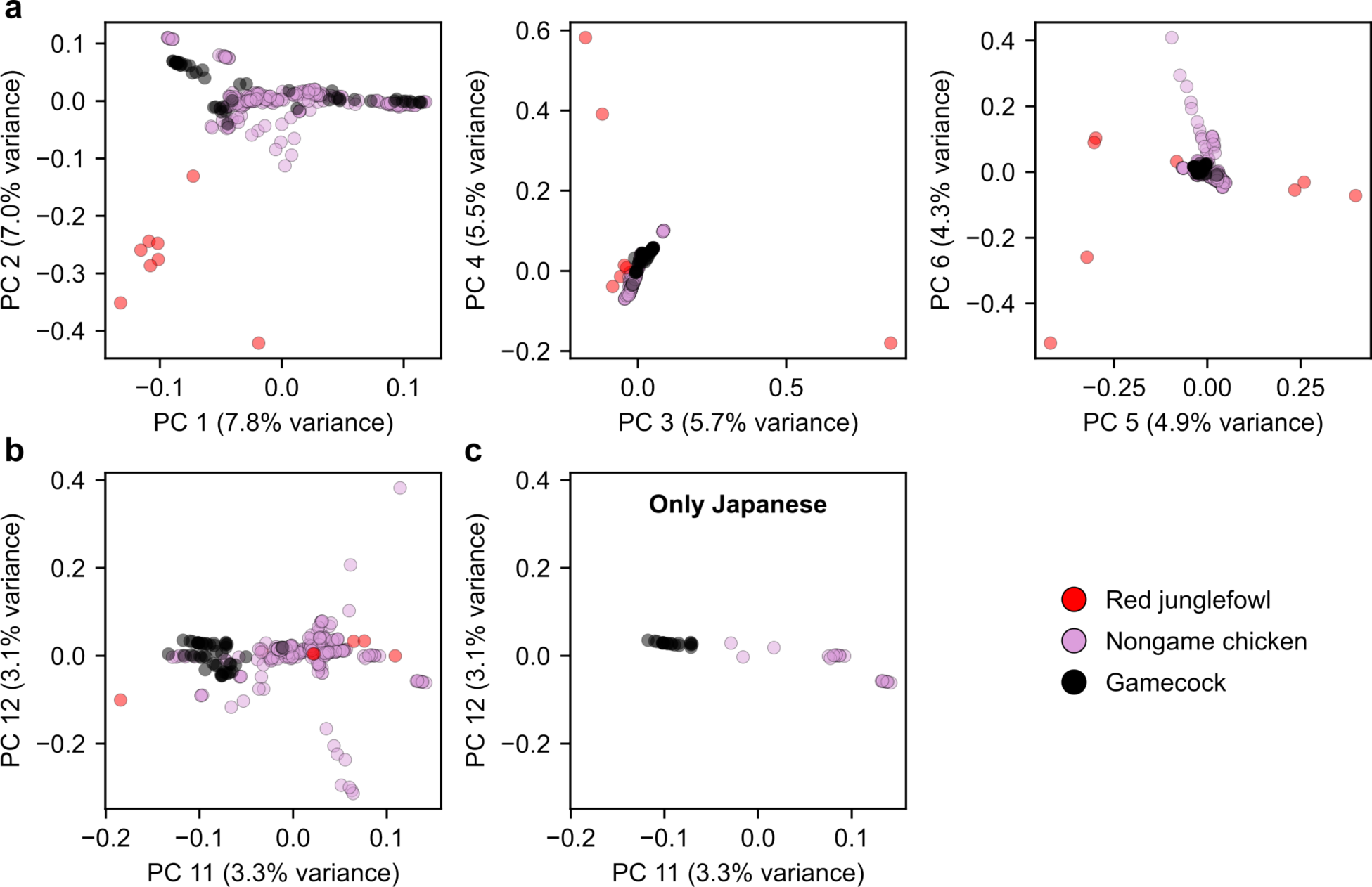
PCA of genetic variation in *Gallus gallus* samples. **a,** PC 1 and 2 separate wild Red junglefowl from domesticated chickens (nongame chickens and gamecocks). **b,** PC 11 mostly separates chickens into nongame chickens and gamecocks. **c,** Same as **b** but including Japanese samples exclusively.

**Suppl. Figure 3.**
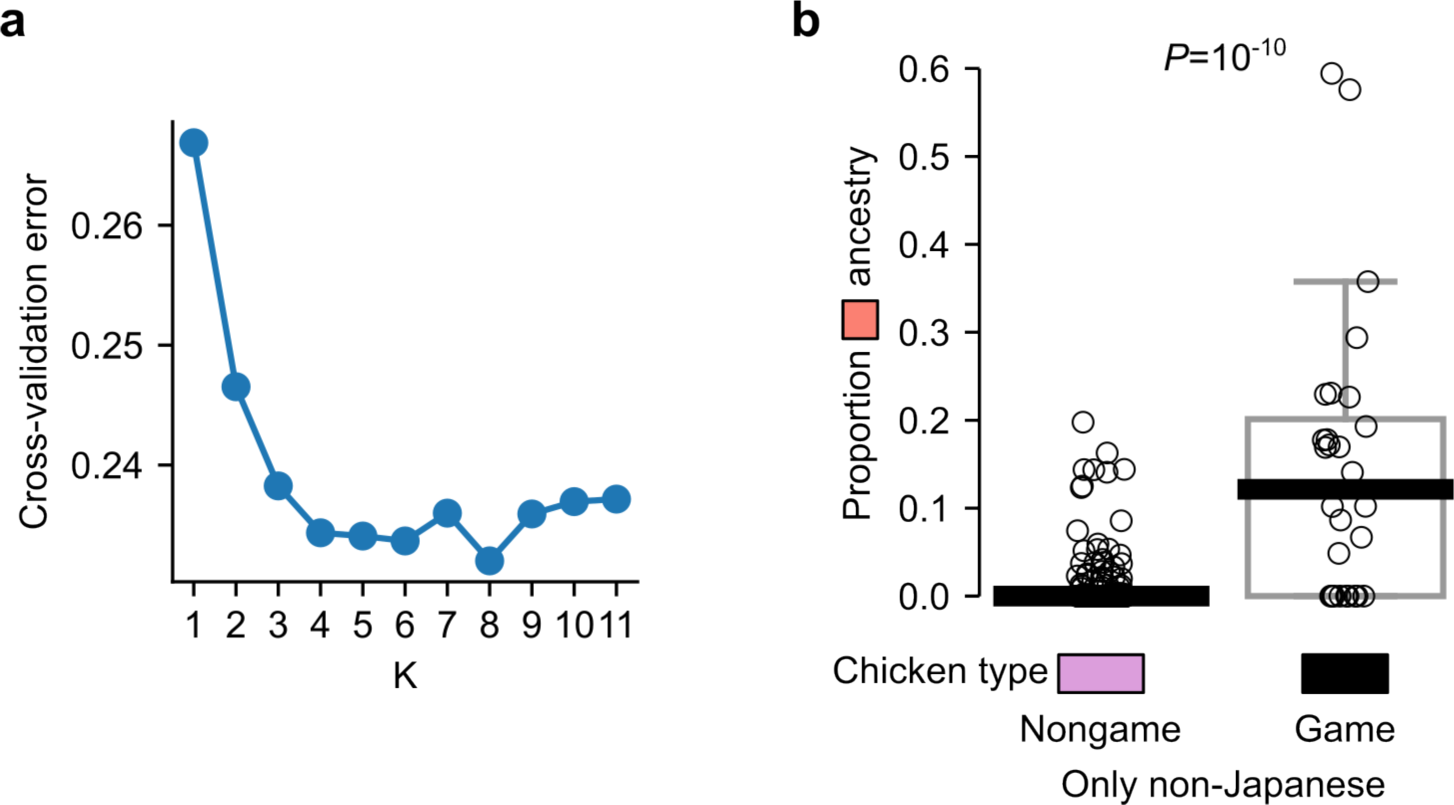
ANCESTRY cross-validation error and analysis without Japanese samples. **a,** Cross-validation error at different values of K. **b,** Proportion "salmon-colored" ancestry in samples that are not from Japan.

**Suppl. Figure 4.**
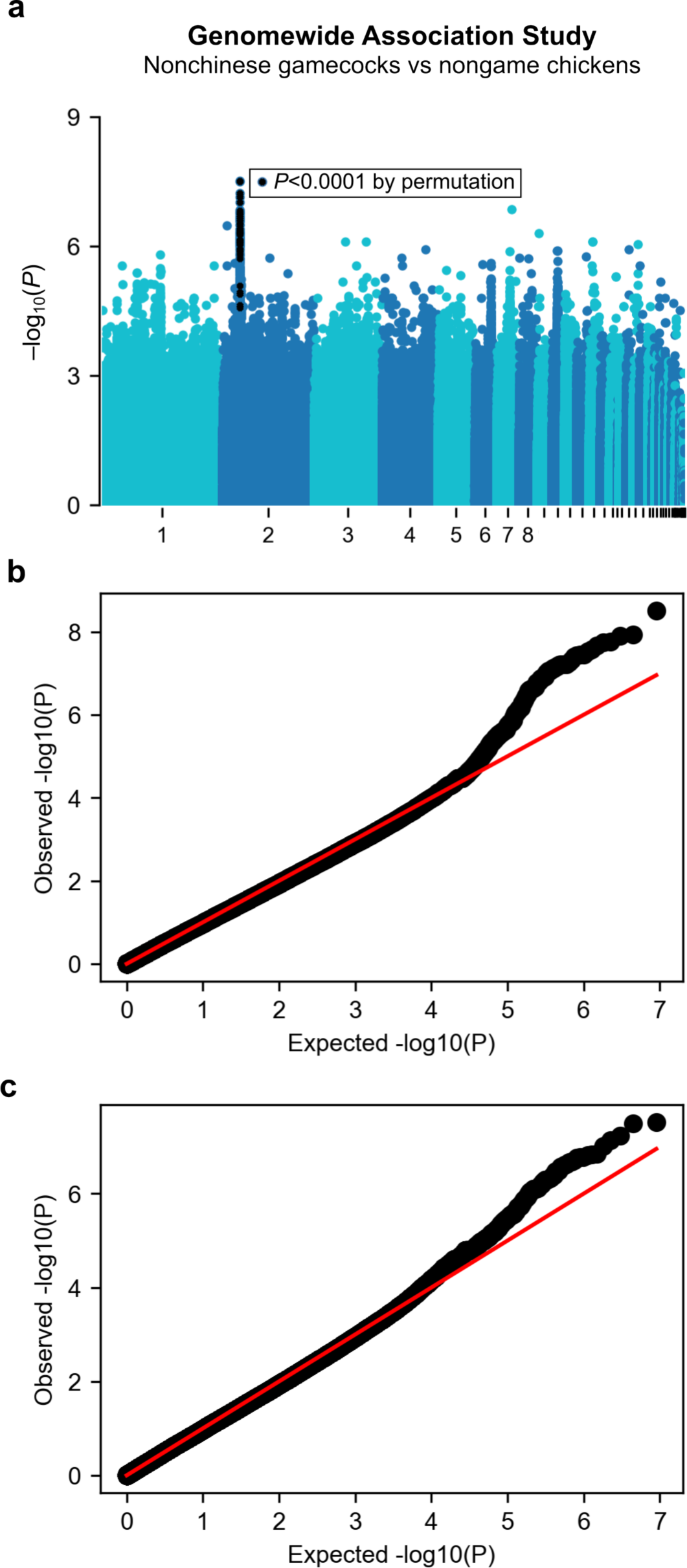
Genome-wide association study excluding Chinese chickens and Q-Q plots. **a,** GWAS of gamecocks that are not from China (n=44) vs nongame chickens (n=62). *P*-values on the y axis using genomic control. Black denotes variants with *P*<10^−4^ by permutation; no variants outside chromosome 2 surpassed that permutation threshold. **b,** Q-Q plot of GWAS in Figure 2a. **c,** Q-Q plot of GWAS in panel **a** of this figure.

**Suppl. Figure 5.**
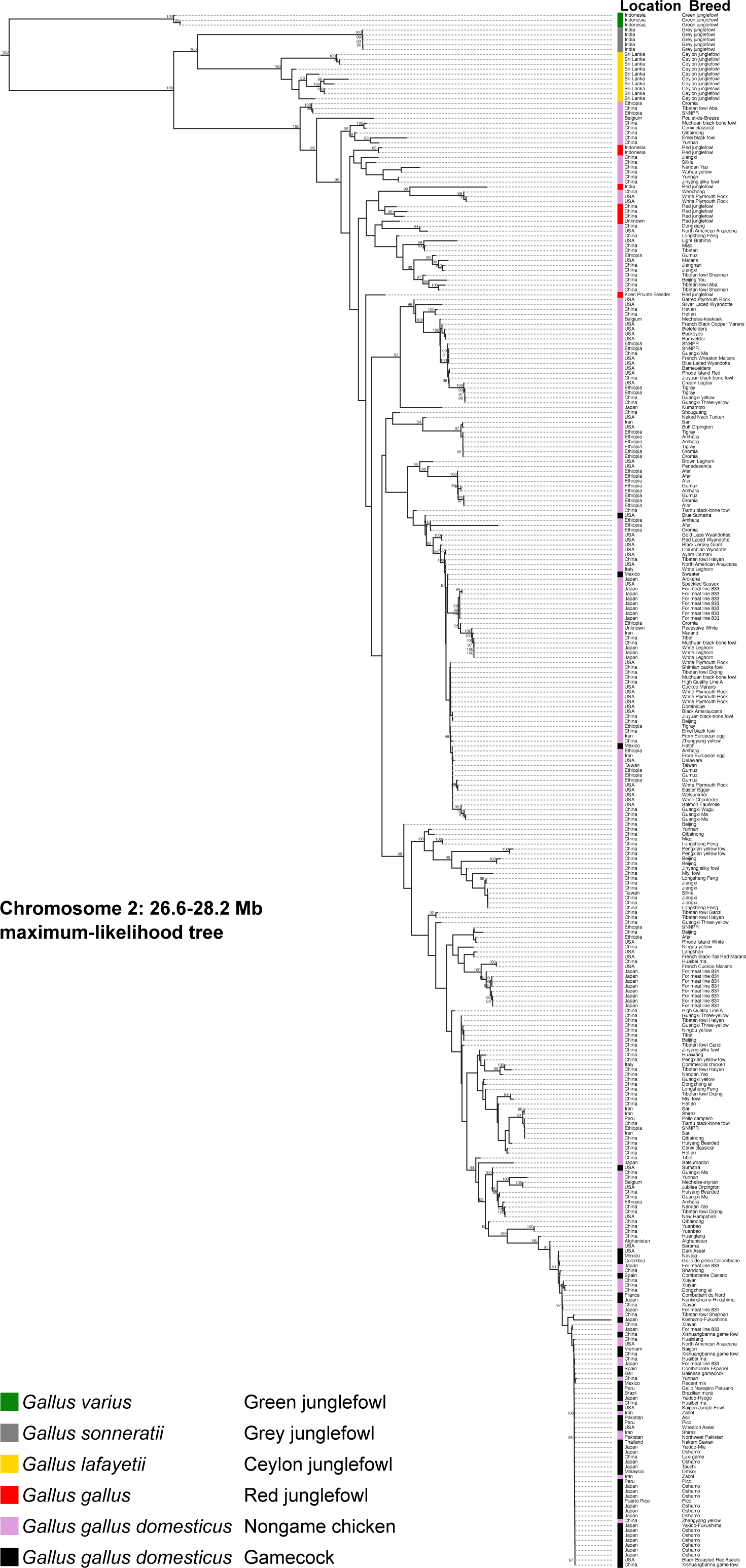
Phylogenetic tree of Chromosome 2 PBS locus that distinguishes gamecocks from nongame chickens. Maximum-likelihood phylogenetic tree of Chromosome 2 locus (26.6–28.2 Mb), including all species in the junglefowl (*Gallus*) genus, as well as gamecocks and nongame chickens from around the world. The name and geographic origin of the samples is presented on the right. Bootstrap support values ≥90 are highlighted on the branches.

**Suppl. Figure 6.**
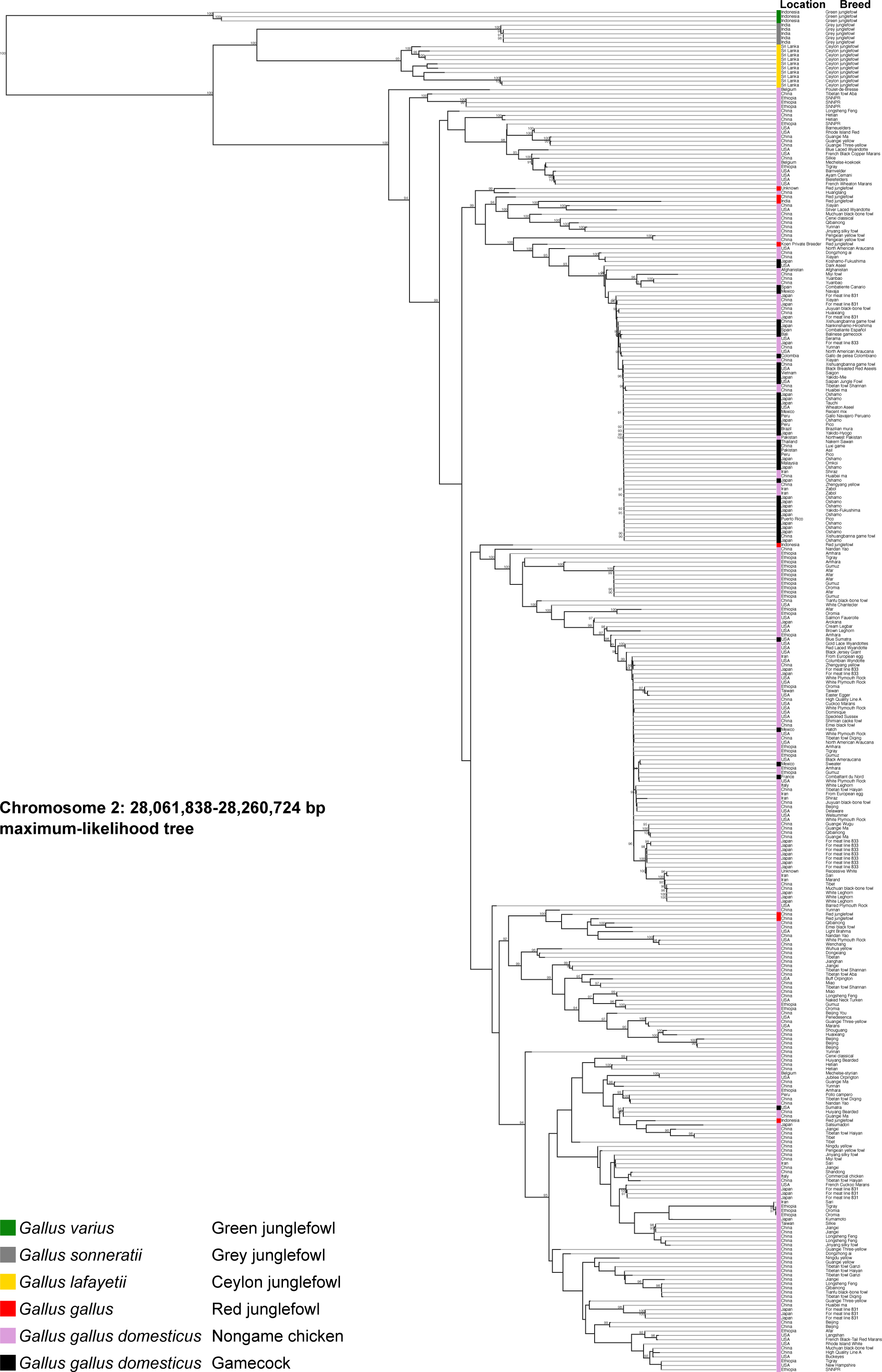
Phylogenetic tree of Chromosome 2 GWAS locus that distinguishes gamecocks from nongame chickens. Maximum-likelihood phylogenetic tree of Chromosome 2 locus (28,061,838–28,260,724 bp), including all species in the junglefowl (*Gallus*) genus, as well as gamecocks and nongame chickens from around the world. The name and geographic origin of the samples is presented on the right. Bootstrap support values ≥90 are highlighted on the branches.

**Suppl. Figure 7.**
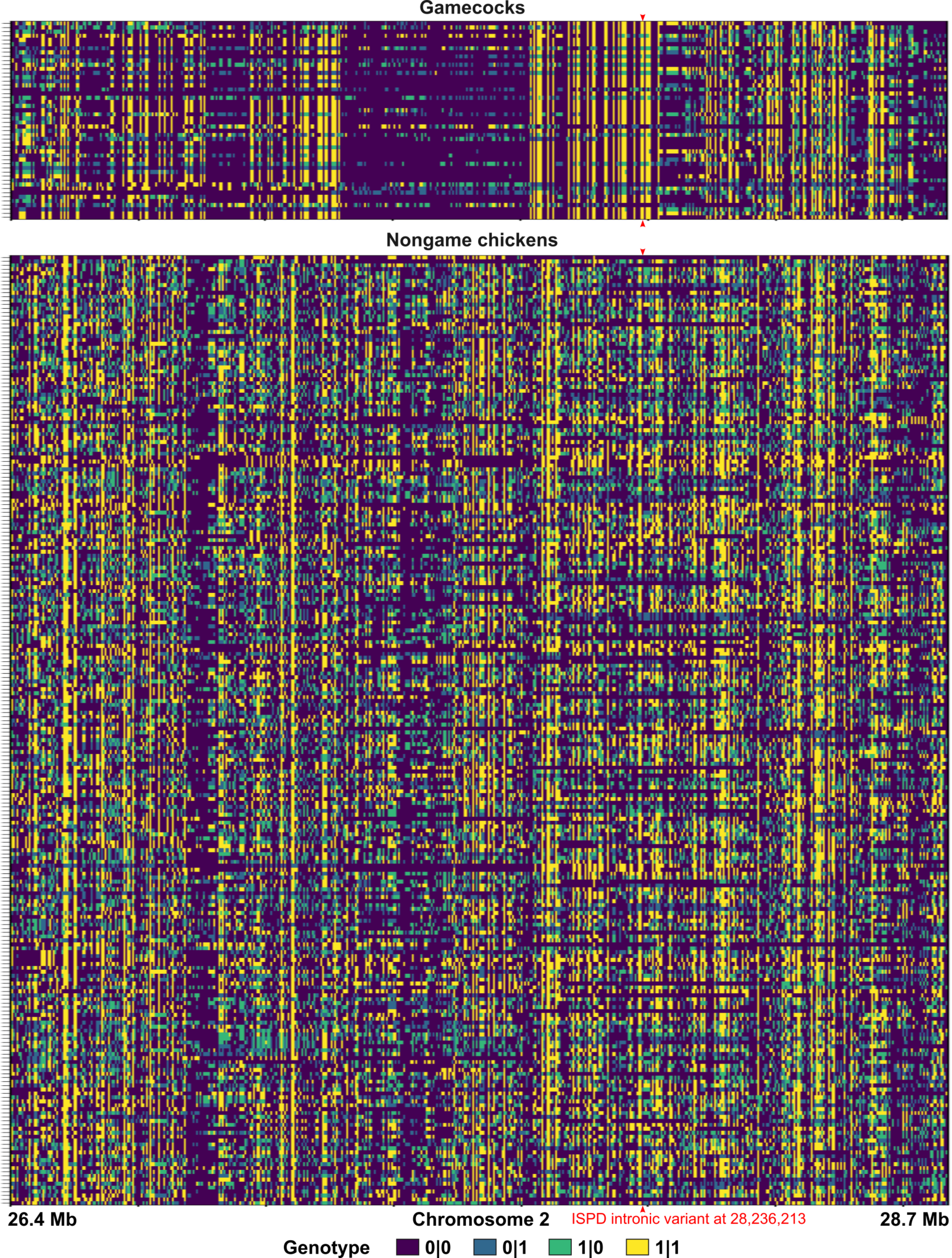
Haplotypes of Chromosome 2: 26.4–28.7 Mb in gamecocks and nongame chickens.

## References

1. Wang, M.-S. et al. 863 genomes reveal the origin and domestication of chicken. Cell Res. 30, 693–701 (2020).

2. Tixier-Boichard, M., Bed’hom, B. & Rognon, X. Chicken domestication: From archeology to genomics. C. R. Biol. 334, 197–204 (2011).

3. Lawler, A. In search of the wild chicken. Science 338, 1020–1024 (2012).

4. Peters, J., Lebrasseur, O., Deng, H. & Larson, G. Holocene cultural history of Red jungle fowl (*Gallus gallus*) and its domestic descendant in East Asia. Quat. Sci. Rev. 142, 102–119 (2016).

5. Lawal, R. A. et al. The wild species genome ancestry of domestic chickens. BMC Biol. 18, 13 (2020).

6. Lawal, R. A. & Hanotte, O. Domestic chicken diversity: Origin, distribution, and adaptation. Anim. Genet. 52, 385–394 (2021).

7. Chickens | Gateway to poultry production and products | Food and Agriculture Organization of the United Nations. https://www.fao.org/poultry-production-products/production/poultry-species/chickens/en/.

8. Lawler, A. Why Did the Chicken Cross the World?: The Epic Saga of the Bird that Powers Civilization. (Atria Books, 2016).

9. Luo, W. et al. Genome diversity of Chinese indigenous chicken and the selective signatures in Chinese gamecock chicken. Sci. Rep. 10, 14532 (2020).

10. Perry-Gal, L., Erlich, A., Gilboa, A. & Bar-Oz, G. Earliest economic exploitation of chicken outside East Asia: evidence from the Hellenistic Southern Levant. Proc. Natl. Acad. Sci. U. S. A. 112, 9849–9854 (2015).

11. Geertz, C. Deep Play: Notes on the Balinese Cockfight. Daedalus 101, 1–37 (1972).

12. Dundes, A. The Cockfight: A Casebook. (University of Wisconsin Press, 1994).

13. Histories of Game Strains (History of Cockfighting Series): Read Country Book. (Read Books Limited, 2013).

14. Bixler, Edsel J. El Gallo Español de Combate. (Editorial Elefante, 2000).

15. Denegri, Marco Aurelio. Arte y Ciencia de la Gallística. (Fondo Editorial, Universidad Inca Garcilaso de la Vega, 2015).

16. Cockfight. *Wikipedia* (2023).

17. Tsudzuki, M. et al. Identification of quantitative trait loci affecting shank length, body weight and carcass weight from the Japanese cockfighting chicken breed, Oh-Shamo (Japanese Large Game). Cytogenet. Genome Res. 117, 288–295 (2007).

18. Herzog, Harold Albert. Studies of the behavior of gamecocks. (University of Tennessee, 1979).

19. McIntyre, R. A. The Game Fowl: Its Origin and History : with Sketches of Great Strains and Their Breeders of Former and Present Times, Together with a Complete Treatise on Breeding and Management of Fowls at Home and at the Pit : Their Diseases and Remedies, and Rules of the Cockpit. (Presses of Grit and Steel, 1906).

20. Cooper, J. W. *Game Fowls*, Their Origin and History, with a Description of the Breeds, Strains, and Crosses: The American and English Modes of Feeding, Training, and Heeling; how to Breed and Cross, Together with a Description and Treatment of All Diseases Incident to Game Fowls. (The Author, 1869).

21. Ulfah, M. et al. Genetic features of red and green junglefowls and relationship with Indonesian native chickens Sumatera and Kedu Hitam. BMC Genomics 17, 320 (2016).

22. Wang, M.-S. et al. Genomic analyses reveal potential independent adaptation to high altitude in Tibetan chickens. Mol. Biol. Evol. 32, 1880–1889 (2015).

23. Fumihito, A. et al. Monophyletic origin and unique dispersal patterns of domestic fowls. Proc. Natl. Acad. Sci. U. S. A. 93, 6792–6795 (1996).

24. Purcell, S. et al. PLINK: A tool set for whole-genome association and population-based linkage analyses. Am. J. Hum. Genet. 81, 559–575 (2007).

25. Nielsen, S. S., et al. Welfare of broilers on farm. EFSA J. 21, e07788 (2023).

26. Guo, L. et al. A missense mutation in ISPD contributes to maintain muscle fiber stability. Poult. Sci. 101, 102143 (2022).

27. Wright, K. M. et al. Dystroglycan organizes axon guidance cue localization and axonal pathfinding. Neuron 76, 931–944 (2012).

28. Picelli, S. et al. Tn5 transposase and tagmentation procedures for massively scaled sequencing projects. Genome Res. 24, 2033–40 (2014).

29. Gheyas, A. A. et al. Integrated environmental and genomic analysis reveals the drivers of local adaptation in African indigenous chickens. Mol. Biol. Evol. 38, 4268–4285 (2021).

30. Fan, W.-L. et al. Genome-wide patterns of genetic variation in two domestic chickens. Genome Biol. Evol. 5, 1376–1392 (2013).

31. Li, D. et al. Genomic data for 78 chickens from 14 populations. GigaScience 6, gix026 (2017).

32. Huang, X. et al. Genome-wide genetic structure and selection signatures for color in 10 traditional Chinese yellow-feathered chicken breeds. BMC Genomics 21, 316 (2020).

33. Noorai, R. E., Shankar, V., Freese, N. H., Gregorski, C. M. & Chapman, S. C. Discovery of genomic variations by whole-genome resequencing of the North American Araucana chicken. PloS One 14, e0225834 (2019).

34. Wang, Y. et al. Multiple ancestral haplotypes harboring regulatory mutations cumulatively contribute to a QTL affecting chicken growth traits. *Commun*. Biol. 3, 1–13 (2020).

35. Guo, Y. et al. A genomic inference of the White Plymouth Rock genealogy. Poult. Sci. 98, 5272–5280 (2019).

36. Guo, Y. et al. Researching on the fine structure and admixture of the worldwide chicken population reveal connections between populations and important events in breeding history. Evol. Appl. 15, 553–564 (2022).

37. Sun, J. et al. Whole-genome sequencing revealed genetic diversity and selection of Guangxi indigenous chickens. PLOS ONE 17, e0250392 (2022).

38. (How to) Map and clean up short read sequence data efficiently – GATK. https://gatk.broadinstitute.org/hc/en-us/articles/360039568932--How-to-Map-and-clean-up-short-read-sequence-data-efficiently.

39. Picard Tools - By Broad Institute. https://broadinstitute.github.io/picard/.

40. Rhie, A. et al. Towards complete and error-free genome assemblies of all vertebrate species. Nature 592, 737–746 (2021).

41. Vasimuddin, Md., Misra, S., Li, H. & Aluru, S. Efficient architecture-aware acceleration of BWA-MEM for multicore systems. in 2019 IEEE International Parallel and Distributed Processing Symposium (IPDPS) 314–324 (2019). doi:10.1109/IPDPS.2019.00041.

42. Danecek, P. et al. Twelve years of SAMtools and BCFtools. GigaScience 10, giab008 (2021).

43. Bergström, A. et al. Insights into human genetic variation and population history from 929 diverse genomes. Science 367, eaay5012 (2020).

44. Heng Li. SNPable Regions. https://lh3lh3.users.sourceforge.net/snpable.shtml.

45. Chen, S., Zhou, Y., Chen, Y. & Gu, J. fastp: an ultra-fast all-in-one FASTQ preprocessor. Bioinformatics 34, i884–i890 (2018).

46. Hanghøj, K., Moltke, I., Andersen, P. A., Manica, A. & Korneliussen, T. S. Fast and accurate relatedness estimation from high-throughput sequencing data in the presence of inbreeding. GigaScience 8, giz034 (2019).

47. Minh, B. Q. et al. IQ-TREE 2: New models and efficient methods for phylogenetic inference in the genomic era. Mol. Biol. Evol. 37, 1530–1534 (2020).

48. Hoang, D. T., Chernomor, O., von Haeseler, A., Minh, B. Q. & Vinh, L. S. UFBoot2: Improving the ultrafast bootstrap approximation. Mol. Biol. Evol. 35, 518–522 (2018).

49. Kalyaanamoorthy, S., Minh, B. Q., Wong, T. K. F., von Haeseler, A. & Jermiin, L. S. ModelFinder: fast model selection for accurate phylogenetic estimates. Nat. Methods 14, 587–589 (2017).

50. Eaton, D. A. R. Toytree: A minimalist tree visualization and manipulation library for Python. Methods Ecol. Evol. 11, 187–191 (2020).

51. Shaun Purcell & Christopher Chang. PLINK 1.9. https://www.cog-genomics.org/plink/1.9/.

52. Chang, C. C. et al. Second-generation PLINK: rising to the challenge of larger and richer datasets. GigaScience 4, 7 (2015).

53. Waskom, M. L. seaborn: statistical data visualization. J. Open Source Softw. 6, 3021 (2021).

54. Alexander, D. H., Novembre, J. & Lange, K. Fast model-based estimation of ancestry in unrelated individuals. Genome Res. 19, 1655–1664 (2009).

55. Yi, X. et al. Sequencing of 50 human exomes reveals adaptation to high altitude. Science 329, 75–78 (2010).

56. Alistair Miles. scikit-allel - Explore and analyse genetic variation — scikit-allel 1.3.3 documentation. https://scikit-allel.readthedocs.io/en/stable/index.html.

57. Korunes, K. L. & Samuk, K. pixy: Unbiased estimation of nucleotide diversity and divergence in the presence of missing data. Mol. Ecol. Resour. 21, 1359–1368 (2021).

58. Samuk, K., Korunes, K., Trevisani, M. D. & Moshiri, N. ksamuk/pixy: pixy 1.2.7.beta1. (2022) doi:10.5281/zenodo.6551490.

59. Hofmeister, R. J., Ribeiro, D. M., Rubinacci, S. & Delaneau, O. Accurate rare variant phasing of whole-genome and whole-exome sequencing data in the UK Biobank. 2022.10.19.512867 Preprint at https://doi.org/10.1101/2022.10.19.512867 (2022).

